# Body mass and latitude as global predictors of vertebrate populations exposure to multiple threats

**DOI:** 10.1101/2022.02.04.479091

**Authors:** Pol Capdevila, Nicola Noviello, Louise McRae, Robin Freeman, Christopher Clements

## Abstract

The interactive effects of multiple threats are one of the main causes of biodiversity loss, yet our understanding of what predisposes species to be impacted by multiple threats remains limited. Here we analyse a global dataset of over 7000 marine, freshwater, and terrestrial vertebrate populations, alongside trait, threat and geographical data, to identify the factors influencing the number of threats a species is subjected to at the population level. Out of a suite of predictors tested, we find that body mass and latitude both are broadly available for vertebrate species, and influence the number of threats a population is subjected to. Larger bodied species and those nearer the equator are typically affected by a higher number of threats. However, whilst this pattern broadly holds across ecosystems for most taxa, amphibians and reptiles show opposing trends. We suggest that latitude and body mass should be considered as key predictors to identify which vertebrate populations are likely to be impacted by multiple threats. These general predictors can help to better understand the impacts of the Anthropocene on global vertebrate biodiversity and design effective conservation policies.

## Introduction

The Anthropocene is characterised by the strong influence of human activities on the structure and functioning of Earth’s natural systems (Steffen *et al.*, 2011; Dirzo *et al.*, 2014). Threats like climate change, habitat loss, exploitation, pollution, or invasive species, directly or indirectly caused by human activities, are reshaping the biosphere at an unprecedented rate and scale (Scholes *et al.*, 2018; Díaz *et al.*, 2019; IPCC, 2021). Although the individual effects of these threats can have strong impacts (Kroeker *et al.*, 2010; Newbold *et al.*, 2015; Hughes *et al.*, 2017), about 80% of species are exposed to more than one threat simultaneously (IUCN 2021). The pervasiveness of multiple threats is of particular concern because of the unpredictability of their interactive effects (Darling & Côté, 2008; Côté *et al.*, 2016). Yet, our understanding of the factors driving exposure to multiple threats remains limited (Maxwell *et al.*, 2016; Hodgson *et al.*, 2017).

Whether a population is exposed to a threat is a result of the combined effects of environmental factors, species life histories, and human activity (Purvis *et al.*, 2000; Cardillo *et al.*, 2005). Life history traits such as body mass, trophic level, or habitat specificity have been linked to the vulnerability of species (Fisher & Owens, 2004; Di Marco *et al.*, 2015; Pacifici *et al.*, 2017). For instance, species with large body mass are disproportionately targeted for exploitation (Pauly *et al.*, 1998; Duncan *et al.*, 2002), making them more vulnerable to further threats. Likewise, predators usually require large home ranges and also depend on the abundance of their prey species, making them vulnerable to habitat loss, as well as being a common target for hunting (Cardillo *et al.*, 2005; Wolf & Ripple, 2016). Moreover, species with low habitat specificity have the potential to be more prone to be exposed to multiple threats, given their wider range of distribution (Malcolm *et al.*, 2006; Ehrlén & Morris, 2015; Batt *et al.*, 2017). While the influence of all these traits on the vulnerability to species extinction has been largely explored (Purvis *et al.*, 2000; Fisher & Owens, 2004; Cardillo *et al.*, 2005), how these contribute to the predisposition of species to being exposed to multiple threats remains unknown.

The exposure of species to threats can also depend on environmental factors. For instance, the prevalence and impact of anthropogenic threats differs in marine, terrestrial and freshwater systems (Díaz *et al.*, 2019). While in freshwater and terrestrial ecosystems habitat loss is the most prevalent threat (Newbold *et al.*, 2015; Birk *et al.*, 2020), exploitation represents the most pressing threat for marine species (Halpern *et al.*, 2015). On top of that, local and global threats show distinct spatial clustering worldwide (Bowler *et al.*, 2020; Harfoot *et al.*, 2021). Many local threats are directly linked to human populations (e.g., habitat loss, hunting, etc.), so their presence is likely to change in line with human population density across different latitudes (Santini *et al.*, 2017). Global threats (e.g. climate change) are also non-uniformly distributed, particularly across latitude (Harfoot *et al.*, 2021; IPCC, 2021), making it challenging to predict them using simple proxies (Sunday *et al.*, 2012).

Understanding the role life history traits and the environmental factors influencing the predisposition of vertebrate populations to be exposed to multiple threats is therefore the first step to manage their effects (Maxwell *et al.*, 2016). Here, we study multiple threats by identifying factors that best predict the number of threats a population is affected by. To do this, we use population-level threat data from the Living Planet Database (Loh *et al.*, 2005), containing spatially explicit data for 7826 populations of 2667 vertebrate species, across the seven continents and all major ecosystems. To test the influence of life history on the predisposition of species to be exposed to multiple threats, we supplemented the threat data with traits which are broadly available and comparable across different taxa: body mass, trophic level, and habitat specificity. To test the influence of environmental factors, we also supplemented the data with human population density, latitude, and system (freshwater, marine or terrestrial) as proxies. We then used multilevel Bayesian models to determine which factors have the strongest influence on the predisposition of populations to be exposed to multiple threats.

## Materials and Methods

### Threats data

To determine the number of threats vertebrate populations are exposed to, we used the Living Planet Database (LPD hereafter). The LPD (http://livingplanetindex.org/data_portal) contains information on over 25,000 vertebrate populations around the world, comprising all vertebrate classes across marine, freshwater, and terrestrial systems and providing population-specific information such as spatial location, abundance, and threat exposure. Data are collected from scientific literature, online databases, and grey literature published since 1970, with at least two years of abundance; detailed inclusion criteria for the LPD can be found in Collen et al., (2009). If the data source was a report of paper, the entire article would be screened and the information was usually extracted from the discussion. For population data shared directly form a data provider, threat information was recorded in the database template form that was provided. A population did not have to be in decline for a threat to be recorded.

Of the 25,054 population time series making up the LPD (including confidential records), 7826 contained data relating to population threat exposure. Based on information from the data source, for each publication we first identified whether the population was threatened, not threatened or whether its threat status was unknown. In this study, we only considered those populations for which threat status information was available. Threats were identified as direct or indirect human activities or processes that impacted the populations for at least 50% of the surveyed years, according to the original source of the time series. If the population was threatened, the number of threats at which the population was exposed was recorded, from one to three. The information within the data sources was sometimes quantitative, e.g. stating number of individuals hunted annually, but most often it was reported in a qualitative way, e.g. a describing a general pattern of hunting that impacts the populations. For this reason, and because the impact of the threat was rarely quantified in the data sources, broad categories describing the threat to the population were recorded.

### Body mass data

Body mass data were collated from a number of pre-existing databases and the scientific literature (see Table S1 for a full list of sources utilised). When minimum and maximum values where given, maximum was taken to ensure measures were most likely those of mature individuals, and thus in line with commonly reported measures from the other databases. Most data sources did not contain sex-specific body mass measurements; however, where sex was indicated an average of the male/female record was taken to account for dimorphism. Finally, where multiple records of the same species were present between datasets, the mean was taken, with all records then standardised to reflect a common unit (g, grams).

For some taxa body mass data were unavailable, and so were estimated using allometric regression equations using length measurements when possible (Feldman *et al.*, 2016; Stark *et al.*, 2020). We used the general equation *W* = *a L*^*b*^, where *W* = body mass, *L* = length, and *a* and *b* are the intercept and slope of a regression line over log◻transformed weight◻at◻length data, respectively (Froese, 2009; Ripple *et al.*, 2017). This method was applied to 47 amphibian species using snout to vent length (SVL) records and clade-specific regression coefficients in FishBase (Froese, 2009; Santini *et al.*, 2018; Stark *et al.*, 2020). A further 320 fish species’ mass were estimated, based on maximum total length (TL) and regression coefficients in FishBase (Froese, 2009). Where a measure other than TL was listed (e.g., standard length (SL), fork length (FL)), regression coefficients were used to convert these to total length before estimating body mass.

### Trophic level data

We broadly classified species according to their diet in three main categories: omnivores, carnivores, or herbivores. For amphibians, birds, mammals and reptiles, we used the data from Etard et al. (2020). For bony and cartilaginous fishes we inferred trophic levels from dietary information obtained from the parameter *Feeding Type* contained in *FishBase* (Froese, 2009). Following the description in Froese (2009) we considered: that herbivores were those species with between 2.0 and 2.19; carnivores had trophic levels equal to or greater than 2.8; and omnivores had trophic levels between 2.2 and 2.79.

### Habitat breadth data

We estimated the habitat breath as the number of distinct habitats a species utilises according to the IUCN habitat classification scheme (Daskalova *et al.*, 2020; Etard *et al.*, 2020). For amphibians, birds, mammals and reptiles, we used the data available in Etard et al. (2020). For bony and cartilaginous fishes the number of habitats was estimated using the *rredlist* package (Chamberlain, 2017).

### Human population density data

To estimate the human influence across different latitudes, we obtained human population density (inhabitants/km^2^) information from HYDE3.2.001 (Hurtt *et al.*, 2011). The human population density represents the number of human habitants per km^2^ per grid cells of 5’ resolution. We used the country where the vertebrate populations were studied to obtain the human population density data of each time series.

### Final dataset

When merging the above datasets with the data from the LPD, not all the species had the same information available. The variables that were accessible for most of the species was latitude (7826 time-series) and body mass (7492), followed by human population density (7361), habitat breadth (6330) and trophic level (4833; Figure 2**a**). When accounting for the combined availability of the variables, trophic level was the variable with the less availability (Figure 2**b**). 1087 populations were missing for the combined factors of trophic level and habitat breadth, 112 for trophic level and body mass, 97 trophic level, body mass and habitat breath, 67 trophic level and human population density, 48 trophic level, human population density and habitat breadth and only 1 for trophic level, human population density and body mass (Figure 2**b**). Because of the low numbers of shared data between some of the factors, we fitted each model (see *Statistical Analysis*) using the dataset with the maximum number of data for each factor. That is, the size of the data set used for each of the models was different depending on the data availability for each of the factors tested.

**Figure 1.**
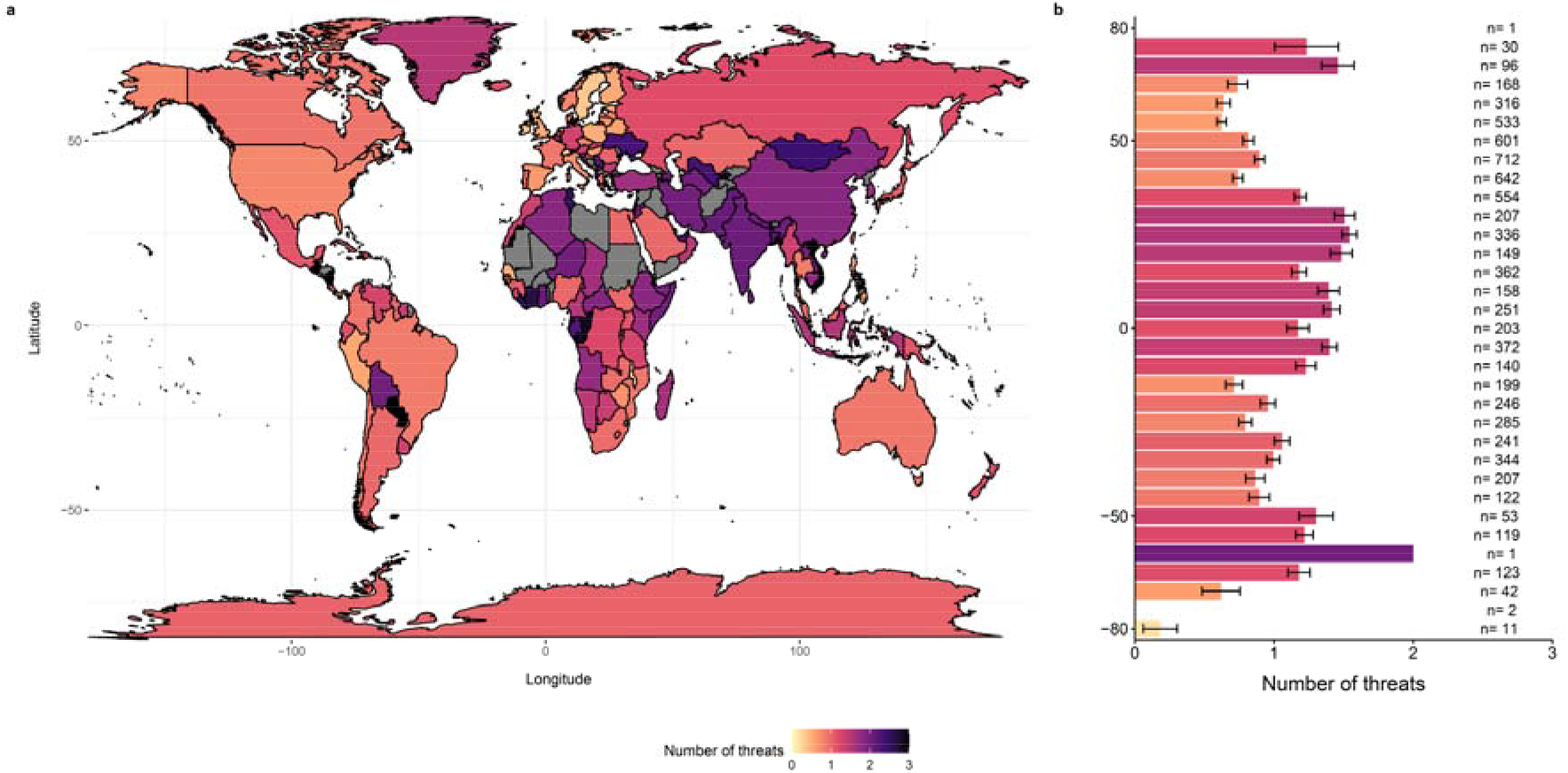
Global distribution of the mean number of threats by country and latitude. Global overview of the mean number of threats, (**a**) within each country and (**b**) by latitude with numbers alongside bars representing sample sizes for each 5° latitude bin.

**Figure 2.**
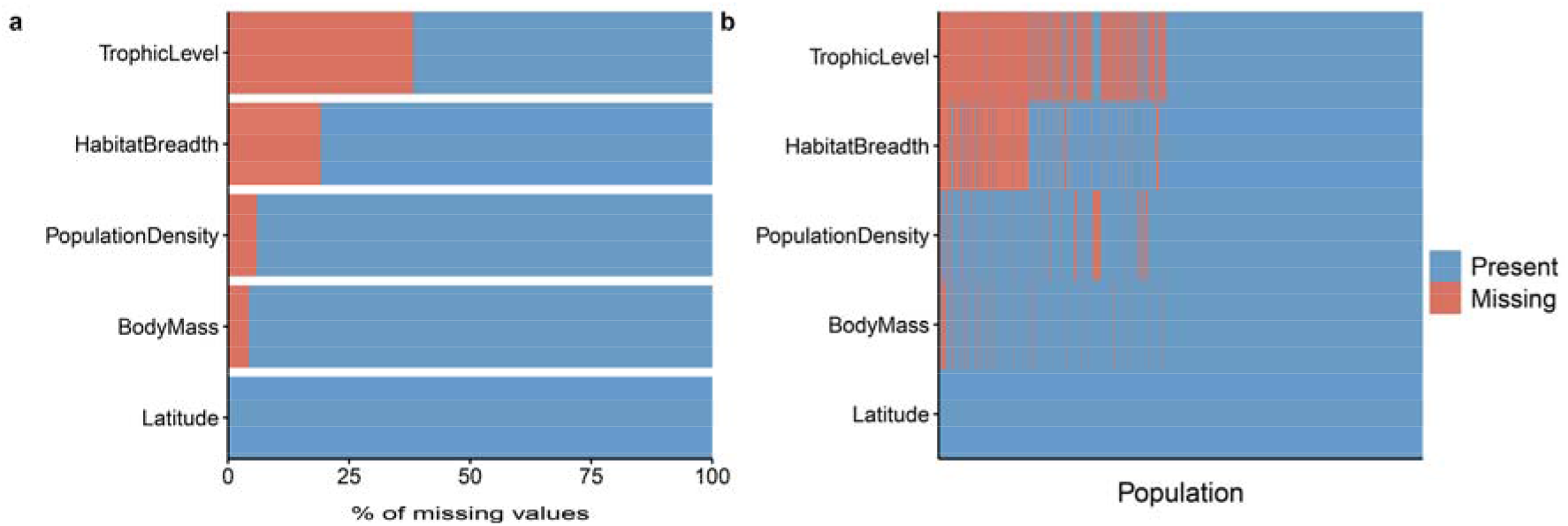
Patterns of missingness in the data. The variable the most available for the species in the subset of data from the Living Planet Database containing threat information was body mass. (**a**) Proportion of missing and present values of the different variables. (**b**) Total presence and absence of the different variables across the dataset.

### Statistical Analysis

To quantify the effects of latitude, body mass, habitat breath, human population density, system, taxon, and trophic level we developed a set of multilevel Bayesian models, using number of threats as a response variable. Body mass was log-transformed and we used the absolute value of latitude. Body mass, latitude, habitat breath, and human population density were added as numeric fixed effects and were all standardised by subtracting the mean from each value and dividing by its standard deviation. System, taxonomic class, and trophic level were considered as categorical variables: the first having three levels, marine, terrestrial, and freshwater; the second having five, amphibians, birds, bony fishes, cartilaginous fishes, mammals and reptiles; and the third having three levels, omnivores, carnivorous and herbivorous.

First, to test the effects of each of the aforementioned factors on the number of threats at which vertebrate populations were exposed we fitted individual models for each of the factors. Then, because we found that the system and taxonomic group had an influence on the number of threats at which populations were exposed (see Results), we fitted individual models for each combination of taxonomic group and system. To account for the non-independence of repeated measurements for each species we included a random intercept for each species. Given the lack of phylogenetic signal in the number of threats at which the populations were exposed (Figure S1) we did not include a phylogeny in these models. Each model with categorical factors (e.g. system, taxon and trophic level) was fitted with a zero intercept to allow us to determine absolute effect of each category of the factors. The general structure of the models was:

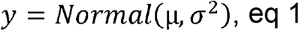

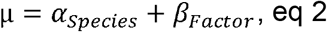

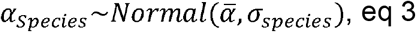

We set weakly informed priors:

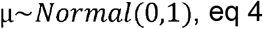

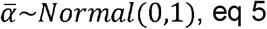

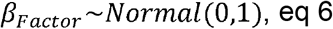

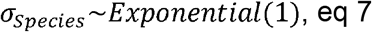

All models were fitted using the brms package v2.1.0 (Bürkner, 2017) in R v4.0.0 (R Core Team, 2020). Models were run for 10000 iterations, with a warmup of 1000 iterations. Convergence was assessed visually by examining trace plots and using Rhat values (the ratio of the effective sample size to the overall number of iterations, with values close to one indicating convergence).

## Results

### General models

The number of threats which populations are exposed to is affected by a number of factors (Figure 3). Among all the systems, freshwater and terrestrial species are exposed to a higher number of threats (Figure 3**a**; Table S3). Reptiles are the taxonomic class exposed to the highest number of threats, followed by amphibians, birds, mammals, cartilaginous fishes, and then bony fishes respectively (Figure 3**b**; Table S3). All trophic levels show similar degree of exposition to multiple threats, with omnivores slightly less at risk than carnivores or herbivores (Figure 3**c**; Table S3). Across all taxa and systems there is low evidence for the influence of body mass on the number of threats (Figure 3**d**; Table S3). On the contrary, latitude has a clear effect on the number of threats, with populations at higher latitudes being exposed to a lower number of threats (Figure 3**e**; Table S3). However, human population densities have a less clear effect, with a low certainty that the effect size is different to zero (Figure 3**f**; Table S3). Finally, there is moderate evidence that species with larger habitat breadth are exposed to a lower number of threats (Figure 3**g**; Table S3).

**Figure 3.**
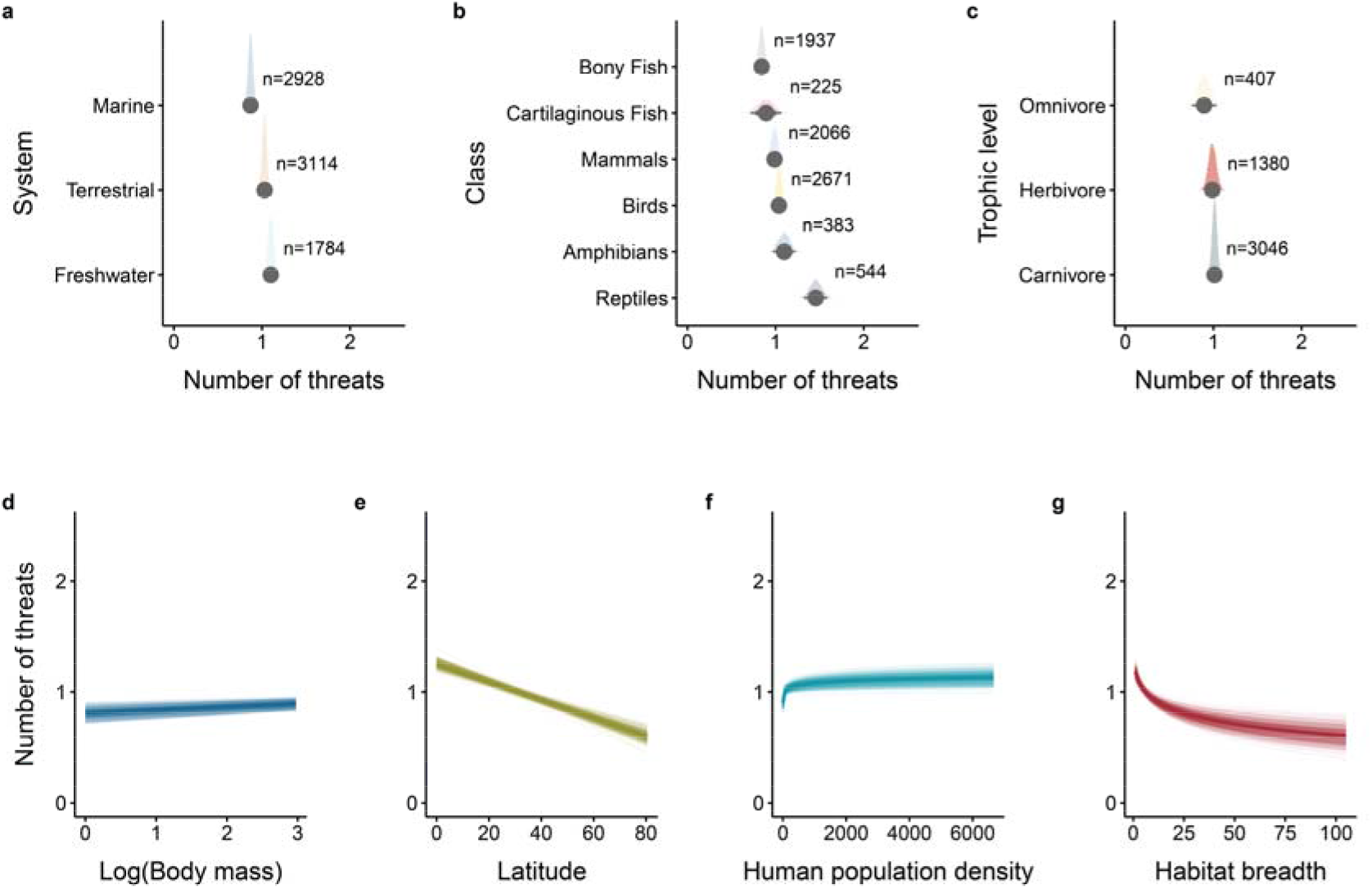
Factors influencing the number of threats at which vertebrate populations are exposed. (**a-c**) density plots of posterior distributions for the effects of (**a**) system, (**b**) taxonomic class, and (**c**) trophic level, on the number of threats. Each density plot is based on 1,000 samples from the posterior distribution of the slope estimates (Table S2). The reported values are the highest posterior density median values (circles), with 80% (thickest bars), 90%, and 95% (thinnest bars) uncertainty intervals. **n**represents the sample size for that given threat in the original dataset. (**d-g**) predictions of the number of threats as a function of the (**d**) body mass (g), (**e**) latitude (absolute value), (**f**) human population density and (**g**) habitat breadth. Lines represent the predictions from the multilevel Bayesian models (Table S2), where thin lines correspond to the predictions drawn from each of the 250 posterior samples of the model, and the thick line represents the mean outcome of the model.

### System and taxa models

The influence of body mass on the number of threats to which populations are exposed varies across different systems and taxa (Figure 4). The number of threats decreases with body mass in amphibians and reptiles independently of the system they inhabit (Figure 4; Table S4). However, these estimates are highly uncertain for freshwater amphibians and reptiles, and marine reptiles (Table S4). For all the other taxonomic groups and systems, the number of threats increases with body mass (Figure 4; Table S4), with high uncertainty in freshwater birds, freshwater and marine bony fishes, and marine mammals (Table S4).

**Figure 4.**
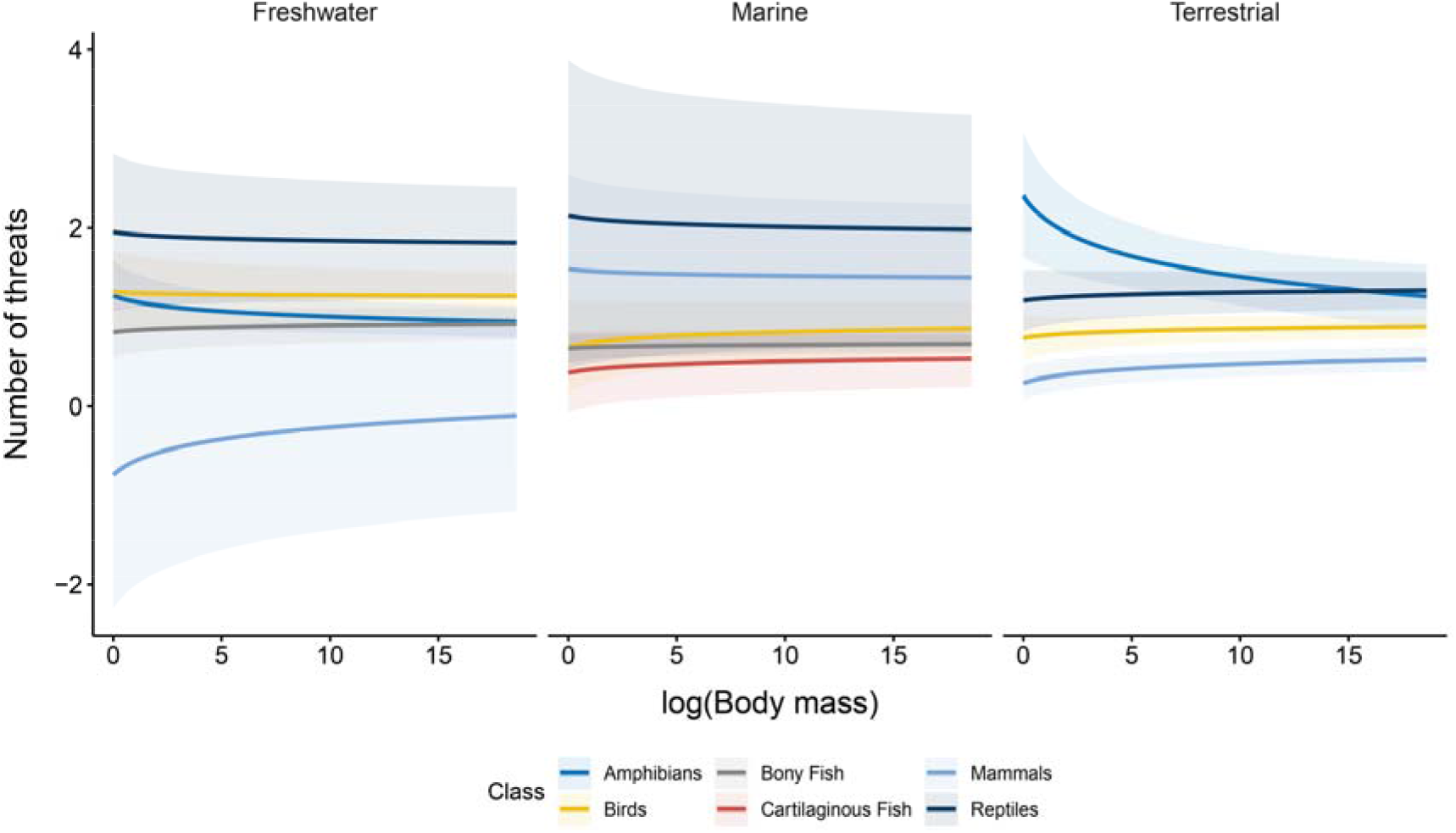
Model predictions of threat number as a function of body mass. Multilevel Bayesian model predictions of the number of threats as a function of body mass (in grams). Ribbons display 95% confidence intervals.

In line with the results of the general models, for most systems and taxa the number of threats decreases at higher latitudes (Figure 5). However, for some system and taxa combinations (notably terrestrial amphibians, birds, marine bony fishes, marine cartilaginous fishes, and marine reptiles) the slope estimates are again uncertain (Table S5). Our results also suggest that the number of threats could increase with latitude in freshwater amphibians, freshwater mammals, and terrestrial reptiles, although again these estimates were highly uncertain (Table S5).

**Figure 5.**
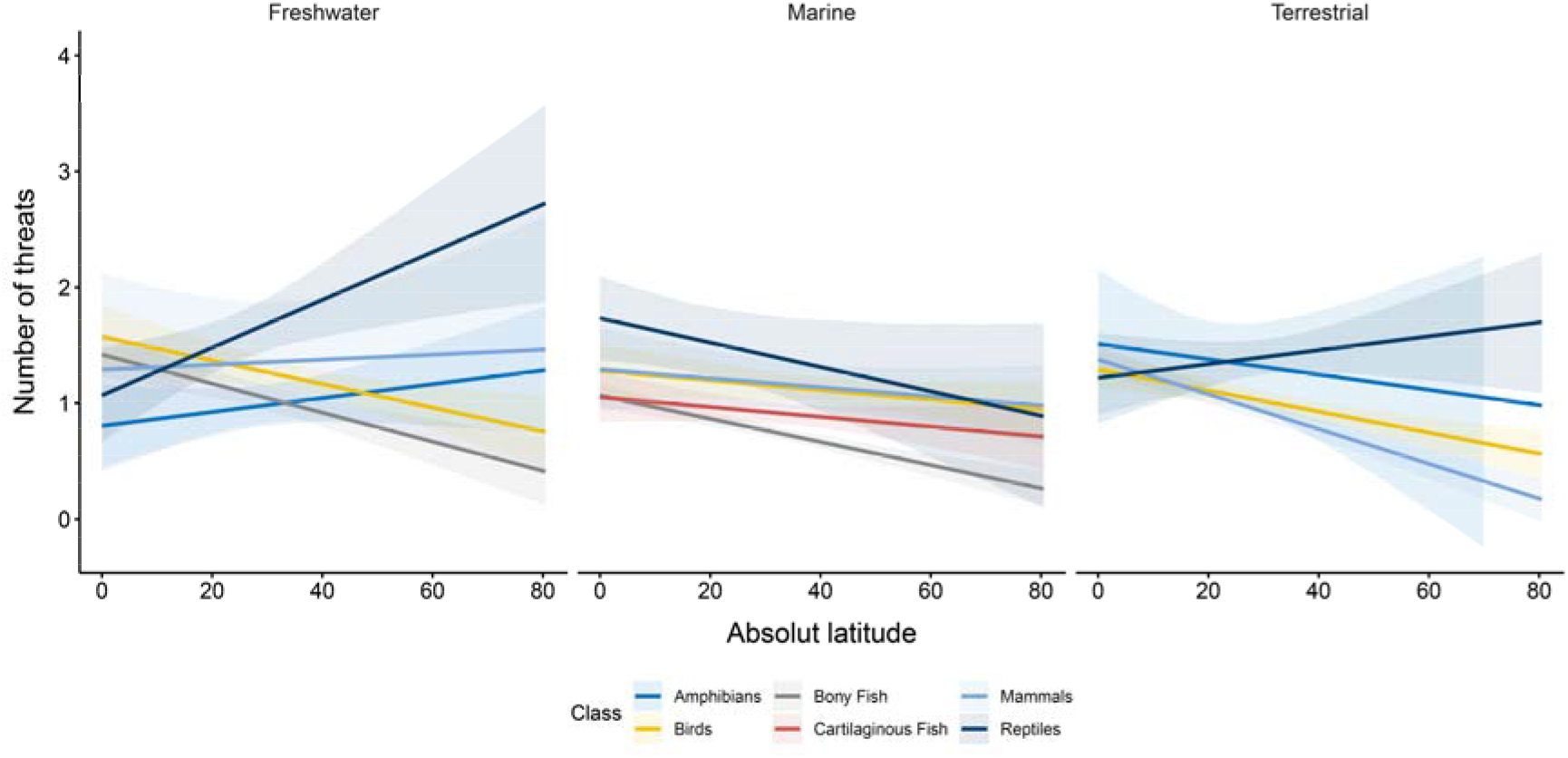
Latitudinal trends of the number of threats. Multilevel Bayesian model predictions of the number of threats as a function of the absolute value of latitude. Ribbons display 95% confidence intervals.

## Discussion

Identifying the factors determining the predisposition of species to be exposed to multiple threats is crucial to maintain biodiversity (Gunderson *et al.*, 2016; Maxwell *et al.*, 2016). To date, most studies have focused on identifying the factors that make species more prone to extinction, rather than to multiple threats (Purvis *et al.*, 2000; Cardillo *et al.*, 2005; Atwood *et al.*, 2020). Consequently, we currently lack understanding of the (a)biotic factors which predispose species to multiple threats, what could help pre-emptively design conservation actions. Here, using a global collation of threat, trait, and geographic data from vertebrate populations, we show that a suite of factors can be used to anticipate the number of threats which vertebrate populations are exposed to. Among these, latitude and body mass are the most readily available and with a strong predictive power. These results are the first necessary step to develop predictive approaches to anticipate the number of threats impacting wildlife populations using minimal data.

Latitude proved a strong predictor of the number of threats which populations are exposed to. Although we hypothesised that the potential reason for this pattern could be that the largest number of people live in lower latitudes (Kummu & Varis, 2011; Figure S4), human population density was a weak driver of threat number. Human population density has long been considered a proxy for anthropogenic disturbance factors (Santini *et al.*, 2017), and arguably the main cause for the ongoing sixth mass extinction event (Ceballos *et al.*, 2020). This premise is based on the assumption that areas with high human density increase the numbers of threats to populations, driving populations beyond the point of recovery (Symes *et al.*, 2018). However, our results suggest that human population density fails to capture the patterns of threats distribution in our global data, and that latitude encapsulates additional latent predictors which predispose populations to be impacted by multiple threats.

Global and local threats are distributed unevenly across the planet (Bowler *et al.*, 2020). For instance, our results suggest that terrestrial and freshwater species are exposed to a higher number of threats compared to marine ones. These findings are in line with the millennia-long human impacts of terrestrial and freshwater systems (McCauley *et al.*, 2015; Van Der Kaars *et al.*, 2017), but may also reflect the difficulty of monitoring species in marine environments. In addition, the presence of threats affecting species can vary within and across countries (Harfoot *et al.*, 2021), often in relation to local governmental conservation policies (Barnes *et al.*, 2016; Amano *et al.*, 2018). Climate change also shows complex spatial patterns, with some mid-latitude regions projected to experience the highest increase in the temperature of the hottest days, while the Arctic is expected to suffer the highest increase in the temperature of the coldest days (IPCC, 2021). In these areas, where the impacts of climate change are likely to become more intense, the interaction with other threats is likely to increase in the coming decades (Bennett et al., 2015). Given the complexity of accounting for such multiple spatial drivers, latitude can provide a simple proxy for multiple threats exposition.

We also show that in most vertebrate groups larger species are exposed to a greater number of threats. The greater vulnerability of larger species is often attributed to different intrinsic and extrinsic factors (Fisher & Owens, 2004; Cardillo *et al.*, 2005). For instance, larger species are disproportionally targeted for exploitation and more affected by invasive species (Bennett & Owens, 1997; Duncan et al., 2002). Also, species with larger body size often occupy higher trophic levels, which are also associated with higher extinction risk (Böhm et al., 2016; Collen et al., 2011). However, our results support recent research (Atwood *et al.*, 2020) showing that species with higher trophic levels are not necessarily exposed to a larger number of threats. Because body size was the most readily available trait, and its tight link with the life history of species (Gaillard *et al.*, 1989), conservation status (Ripple *et al.*, 2017) and ecological processes (White *et al.*, 2007), our findings validate its use as a proxy for multiple threats exposition.

Amphibians and reptiles were the exception to the abovementioned pattern, with body size being inversely related with number of threats. These pattern may be driven by the nature of the threats affecting them. Both groups are mostly affected by habitat loss, while mammals, birds and fishes are mostly impacted by exploitation (Díaz *et al.*, 2019; Harfoot *et al.*, 2021). While larger individuals are often the target of exploitation (Pauly *et al.*, 1998; Duncan *et al.*, 2002), the lower dispersal ability and more constrained range sizes of small organisms could make them more vulnerable to habitat loss (Cardillo *et al.*, 2008; González-Suárez *et al.*, 2013; Pacifici *et al.*, 2017). Moreover, our results suggest amphibians and reptiles are facing the largest number of threats, mirroring recent reports suggesting that these are the vertebrate groups experiencing the most dramatic decline (Daskalova *et al.*, 2020). Despite these findings, amphibians and reptiles are the most understudied vertebrate groups in global conservation assessments (Alroy, 2015). For instance, about 25% of known reptiles and amphibian species are classified as data deficient by the IUCN Red List (IUCN, 2021). Hence, our results adds evidence for the need of global efforts to study these groups, to better understand the causes of their decline and develop effective conservation policies (Gibbons *et al.*, 2000). Our findings also highlight the importance of understanding the mechanisms that predispose reptiles and amphibians to multiple threats as a key area for future research.

Although our research uses the largest compilation of population-level threat data, there are still gaps in our understanding of the drivers of multiple threats. While the LPD draws from published literature, this also means its data inherits any biases derived from its sources. This has resulted in the over-representation of well-studied regions and taxa, with research also inclined towards populations within protected areas and terrestrial ecosystems (Loh *et al.*, 2005; McRae *et al.*, 2017). In addition, while here we only focused on the number of threats, their type (e.g. exploitation, habitat loss), intensity and/or frequency also has a major influence on the population trends and this information was not readily available. For instance, different threats, or the same threat with different intensity and/or frequency, might interact in different ways, causing different impacts on the populations (Darling & Côté, 2008; Côté *et al.*, 2016; Orr *et al.*, 2020). Not only that, but the timing (when threats impacted the population over the time series) and the synchrony (the overlap on time between multiple threats) of the threats might have a strong influence on populations (Johnstone *et al.*, 2016; Jackson *et al.*, 2021). The limited data available on disturbances nature and timing at the population-level hampered including such information in our analyses. To this end, we advocate explicit reference to threats within ecological research to enable the expansion of current databases and to keep multiple threats processes at the forefront of developing research.

