## Supplementary material for "Body mass and latitude as global predictors of vertebrate populations exposure to multiple threats": Table S1, Table S2, Table S3, Table S4, Table S5

COntent

### Supplementary Data

**Table S1. Body mass and length data sources.** This table includes all sources of body mass data used within the study, plus measurements used in allometric mass/length equation calculations.

| **Data** | **Measurement** | **Reference** |
| --- | --- | --- |
| **Amniote** | Body Mass | Nathan P. Myhrvold, Elita Baldridge, Benjamin Chan, Dhileep Sivam, Daniel L. Freeman, S. K. Morgan Ernest. 2015. An amniote life-history database to perform comparative analyses with birds, mammals, and reptiles. Ecology 96: 3109 |
| **AmphiBIO** | Body Mass | Oliveira, B., São-Pedro, V., Santos-Barrera, G. et al. AmphiBIO, a global database for amphibian ecological traits. Sci Data 4, 170123 (2017) |
| ***Atelopus longirostris*** | Body mass | Elicio Eladio Tapia, Luis Aurelio Coloma, Gustavo Pazmiño-Otamendi & Nicolás Peñafiel (2017) Rediscovery of the nearly extinct longnose harlequin frog Atelopus longirostris (Bufonidae) in Junín, Imbabura, Ecuador, Neotropical Biodiversity,  3: 1, 157-167, DOI:  10.1080/23766808.2017.1327000 |
| ***Chalcorana (Rana) chalconota*** | SVL | Robert F. Inger, Bryan L. Stuart, Djoko T. Iskandar, Systematics of a widespread Southeast Asian frog, Rana chalconota (Amphibia: Anura: Ranidae), Zoological Journal of the Linnean Society, Volume 155, Issue 1, January 2009, Pages 123–147, https: //doi.org/10.1111/j.1096-3642.2008.00440.x |
| **Elton Traits** | Body Mass | Smith et al 2003, Dunning 2007 – see Elton traits metadata |
| **Encyclopedia of Life** | Body Mass | Parr, C. S., N. Wilson, P. Leary, K. S. Schulz, K. Lans, L. Walley, J. A. Hammock, A. Goddard, J. Rice, M. Studer, J. T. G. Holmes, and R. J. Corrigan, Jr. 2014. The Encyclopedia of Life v2: Providing Global Access to Knowledge About Life on Earth. Biodiversity Data Journal 2: e1079, doi:10.3897/BDJ.2.e1079 |
| **Fishbase** | Length (TL / FL / SL) | Froese R. & Pauly D. (eds). (2020). FishBase (version Feb 2018). In: Species 2000 & ITIS Catalogue of Life, 2020-09-01 Beta (Roskov Y.; Ower G.; Orrell T.; Nicolson D.; Bailly N.; Kirk P.M.; Bourgoin T.; DeWalt R.E.; Decock W.; Nieukerken E. van; Penev L.; eds.). Digital resource at www.catalogueoflife.org/col. Species 2000: Naturalis, Leiden, the Netherlands. ISSN 2405-8858. |
| **Handbook of the Birds of the World Alive** | Body Mass | S. M. Billerman, B. K. Keeney, P. G. Rodewald, and T. S. Schulenberg (Editors) (2020). Birds of the World. Cornell Laboratory of Ornithology, Ithaca, NY, USA. https://birdsoftheworld.org/bow/home |
| ***Leiopelma archeyi*** | Body mass | Stark, G,  Meiri, S.  Cold and dark captivity: Drivers of amphibian longevity. Global Ecol Biogeogr.   2018; 27:  1384– 1397. https: //doi.org/10.1111/geb.12804 |
| ***Litoria australis* aka *Ranoidea australis, Litoria dahlia* aka *Ranoidea dahlii, Ranoidea genimaculata* aka *Litoria genimaculata*** | SVL | Vanderduys, E. (2019). Field Guide to the Frogs of Queensland. In Field Guide to the Frogs of Queensland. doi: 10.1071/9780643108790 |
| ***Litoria nannotis* aka *Ranoidea nannotis*** | Body mass | Liem, D.S. (1974). A review of the Litoria nannotis species group and a description of a new species of Litoria from north-east Queensland, Australia. Memoirs of the Queensland Museum 17(1), 151-168.  Cogger, H.G. (1994). Reptiles and Amphibians of Australia. Reed Books, Sydney.  McDonald, K.R. & Alford, R.A. (1999). A Review of Declining Frogs in Northern Queensland. Pp 14-22 in A. Campbell (ed), Declines and Disappearances of Australian Frogs. Environment Australia, Canberra. 234 pp. |
| ***Myotis escalerai*** | Body mass | Quetglas, J. (2016). Murciélago ratonero ibérico – Myotis escalerai. En:  Enciclopedia Virtual de los Vertebrados Españoles. Salvador, A., Barja, I. (Eds.). Museo Nacional de Ciencias Naturales, Madrid. |
| **PanTHERIA** | Body Mass | Kate E. Jones, Jon Bielby, Marcel Cardillo, Susanne A. Fritz, Justin O'Dell, C. David L. Orme, Kamran Safi, Wes Sechrest, Elizabeth H. Boakes, Chris Carbone, Christina Connolly, Michael J. Cutts, Janine K. Foster, Richard Grenyer, Michael Habib, Christopher A. Plaster, Samantha A. Price, Elizabeth A. Rigby, Janna Rist, Amber Teacher, Olaf R. P. Bininda-Emonds, John L. Gittleman, Georgina M. Mace, and Andy Purvis. 2009. PanTHERIA: a species-level database of life history, ecology, and geography of extant and recently extinct mammals. Ecology 90: 2648. |
| ***Rana tavasensis*** | SVL | Düşen, S. (2012). First data on the helminth fauna of a locally distributed mountain frog, “Tavas frog” Rana tavasensis Baran & Atatür, 1986 (Anura: Ranidae), from the inner-west Anatolian region of Turkey. Turkish Journal of Zoology, 36, 496-502. |
| ***Trachycephalus venulosus*** | Body mass | Domingos J. Rodrigues, Masao Uetanabaro & Frederico S. Lopes (2005) Reproductive patterns of Trachycephalus  venulosus (Laurenti, 1768) and Scinax fuscovarius (Lutz, 1925) from the Cerrado, Central Brazil, Journal of Natural  History, 39: 35, 3217-3226, DOI:  10.1080/00222930500312244 |
| **Various amphibian** | Body Mass | Santini L., Benítez-López A., Ficetola G.F., Huijbregts M.A.J. 2017. Length – Mass allometries in Amphibians. Integrative Zoology, 13: 36-45. doi:10.1111/1749-4877.12268 |
| **Various amphibian** | Body mass | Stark, G, Pincheira‐Donoso, D,  Meiri, S.  No evidence for the ‘rate‐of‐living’ theory across the tetrapod tree of life.  Global Ecol Biogeogr.  2020; 00:  1– 28. https: //doi.org/10.1111/geb.13069 |
| **Various amphibian** | SVL | AmphibiaWeb. 2020. <https://amphibiaweb.org> University of California, Berkeley, CA, USA. |
| **Various amphibians** | Body Mass | Trochet A, Moulherat S, Calvez O, Stevens V, Clobert J, Schmeller D (2014) A database of life-history traits of European amphibians. Biodiversity Data Journal 2: e4123. |
| **Various avian** | Body Mass | Terje Lislevand, Jordi Figuerola, and Tamás Székely. 2007. Avian body sizes in relation to fecundity, mating system, display behavior, and resource sharing. Ecology 88: 1605 |
| **Various avian** | Body Mass | Renner, S.C., Hoesel, W. Ecological and Functional Traits in 99 Bird Species over a Large-Scale Gradient in Germany. Data 2017, 2, 12. |
| **Various mammals** | Body Mass | Smith, F.A., Lyons, S.K., Ernest, S.K.M., Jones, K.E., Kaufman, D.M., Dayan, T., Marquet, P.A., Brown, J.H. and Haskell, J.P. (2003), Body mass of late quaternary mammals. Ecology, 84: 3403-3403. |
| **Various primates** | Body Mass | Galán-Acedo, C., Arroyo-Rodríguez, V., Andresen, E. et al. Ecological traits of the world’s primates. Sci Data 6, 55 (2019) doi: 10.1038/s41597-019-0059-9 |
| **Various vertebrates** | Body Mass | Anthony I. Dell, Samraat Pawar, Van M. Savage. 2013. The thermal dependence of biological traits. Ecology 94: 1205. |

### Supplementary analyses

#### Phylogenetic signal

To explore the role of the phylogenetic relatedness of the species included in our analyses in the patterns of exposure to multiple threats, we constructed species-level phylogenetic trees for each of the major taxonomic groups. We obtained the phylogenies for amphibians, birds, mammals and reptiles from https://vertlife.org. For each of the taxonomic groups we downloaded 100 trees and randomly chose 1 for the analysis. Species-level phylogenies for cartilaginous and bony fishes haven’t been resolved yet, so we used data from the Open Tree of Life (OTL, https://tree.opentreeoflife.org). The OTL combines publicly available taxonomic and phylogenetic information across the tree of life (Hinchliff et al., 2015). For bony and cartilaginous fishes we built separate taxonomic trees using the *rotl* R package (Michonneau et al., 2016). Then, we used the package *date life* to account for the phylogenetic distance of the species included (Sánchez-Reyes & O’Meara, 2019). Polytomies (i.e., a node in the tree with >2 species with a common immediate ancestor) were resolved using the *multi2di* function from the *ape* package (Paradis et al., 2004).

To explore the influence of phylogenetic relatedness on the predisposition to be exposed to multiple threats, we fitted hierarchical Bayesian models (see Statistical analyses in the main text). We used the number of threats as response variable but without fixed effects and including the variance-covariance matrix of the phylogeny and species as a random effects. We assessed the phylogenetic signal by calculating the explained variance of the random effects (phylogeny and species) in the posterior distributions of the models. A distribution pushed up against 0 indicates a lack of phylogenetic signal, given that the variance explained is bounded by 0 and only has positive values.

The general structure of the model was:

$y=Normal\left( \mu, \sigma^{2} \right)$, eq 1

$\mu=\alpha_{Species}+\gamma_{Phylogeny}$, eq 2

$\alpha_{Species}\sim Normal(\bar{\alpha}, \sigma_{species})$, eq 3

$\gamma_{Phylogeny}\sim MVNormal\left( 0, \boldsymbol{S} \right)$, eq 4

$\boldsymbol{S}$=$\sigma_{Phylogeny}^{2}$**V**, eq 5

We set weakly informed priors:

$\mu\sim Normal(0,1)$, eq 6

$\bar{\alpha}\sim Normal(0,1)$, eq 7

$\sigma_{Phylogeny}^{2}\sim Exponential(1)$, eq 8

$\sigma_{Species}\sim Exponential(1)$, eq 9

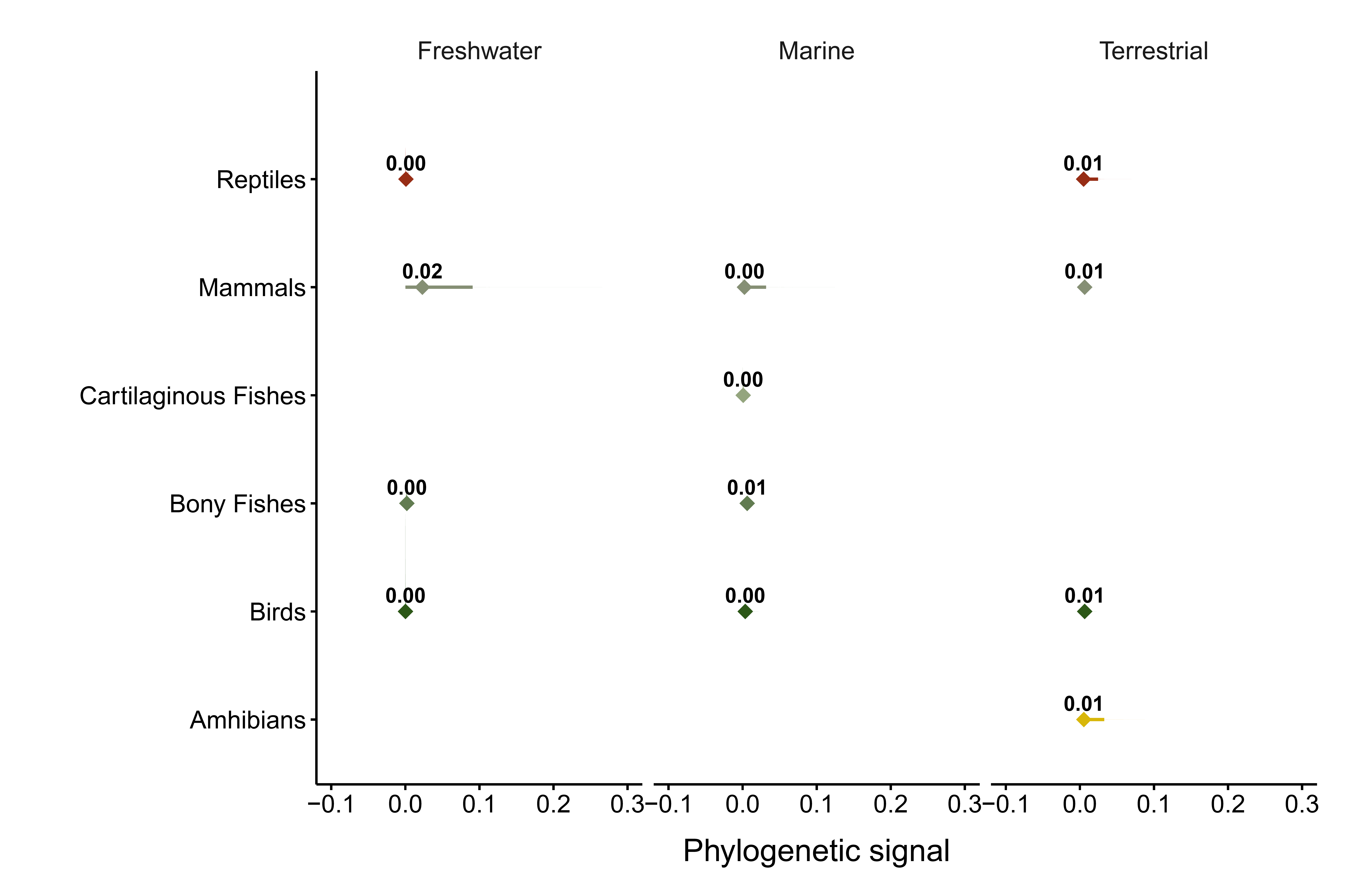
**Fig. S1. Posterior distribution for the phylogenetic signal of the number of threats across the different taxonomic class and system.** The distributions represent the influence of the phylogenetic relatedness in the patterns of variation of the number of threats observed among species. A distribution pushed up against 0 indicates a lack of phylogenetic signal, given that the variance explained is bounded by 0 and only has positive values. The lines below the distribution represent the 95% confidence intervals, with the dot representing the median value.

**Table S2. Outputs of the general models.** Median represents the median of the posterior distribution. CI low and high are the lower and higher values of the 95% confidence interval. ROPE represents the region of practical equivalence, which defines an area (lower and high ROPE) where the value of the parameters is very likely to have a negligible effect (Kruschke, 2014; McElreath, 2020). %ROPE represents the proportion of the 89% Highest Density Interval falling within the ROPE area. If all the HDI falls within the ROPE area (100%) the effects of the parameters can be considered negligible (Kruschke, 2014; McElreath, 2020). The closer the proportion inside the ROPE is to zero, the more confident are the effects of the parameters. Rhat is the ratio of the effective sample size to the overall number of iterations, with values close to one indicating convergence values.

| Model | Parameter | Median | CI (low) | CI (high) | ROPE (low) | ROPE (high) | %ROPE | Rhat |
| --- | --- | --- | --- | --- | --- | --- | --- | --- |
| System | Freshwater | 1.11 | 1.05 | 1.17 | -0.08 | 0.08 | 0 | 1.00 |
|  | Marine | 0.84 | 0.79 | 0.89 | -0.08 | 0.08 | 0 | 1.00 |
|  | Terrestrial | 1.04 | 1.00 | 1.10 | -0.08 | 0.08 | 0 | 1.00 |
| Class | Amphibians | 1.11 | 0.98 | 1.24 | -0.08 | 0.08 | 0 | 1.00 |
|  | Birds | 1.04 | 0.98 | 1.09 | -0.08 | 0.08 | 0 | 1.00 |
|  | Bony Fish | 0.84 | 0.79 | 0.90 | -0.08 | 0.08 | 0 | 1.00 |
|  | Cartilaginous Fish | 0.89 | 0.71 | 1.05 | -0.08 | 0.08 | 0 | 1.00 |
|  | Mammals | 0.99 | 0.91 | 1.06 | -0.08 | 0.08 | 0 | 1.00 |
|  | Reptiles | 1.45 | 1.32 | 1.59 | -0.08 | 0.08 | 0 | 1.00 |
| Trophic level | Carnivore | 1.00 | 0.95 | 1.06 | -0.09 | 0.09 | 0 | 1.00 |
|  | Herbivore | 0.99 | 0.90 | 1.08 | -0.09 | 0.09 | 0 | 1.00 |
|  | Omnivore | 0.92 | 0.78 | 1.06 | -0.09 | 0.09 | 0 | 1.00 |
| Body mass | Intercept | 1.04 | 1.00 | 1.07 | -0.08 | 0.08 | 0 | 1.00 |
|  | Body mass | 0.07 | 0.04 | 0.11 | -0.08 | 0.08 | 76 | 1.00 |
| Latitude | Intercept | 0.89 | 0.86 | 0.93 | -0.08 | 0.08 | 0 | 1.00 |
|  | Latitude | -0.18 | -0.20 | -0.16 | -0.08 | 0.08 | 0 | 1.00 |
| Human population density | Intercept | 0.98 | 0.95 | 1.02 | -0.08 | 0.08 | 0 | 1.00 |
|  | Human population density | 0.05 | 0.04 | 0.06 | -0.08 | 0.08 | 100 | 1.00 |
| Habitat breadth | Intercept | 1.01 | 0.97 | 1.05 | -0.09 | 0.09 | 0 | 1.00 |
|  | Habitat breadth | -0.07 | -0.11 | -0.04 | -0.09 | 0.09 | 77 | 1.00 |

**Table S3. Outputs of the specific body mass models.** Median represents the median of the posterior distribution. CI low and high are the lower and higher values of the 95% confidence interval. ROPE represents the region of practical equivalence, which defines an area (lower and high 95% ROPE) where the value of the parameters is very likely to have a negligible effect (Kruschke, 2014; McElreath, 2020). %ROPE represents the proportion of the 89% Highest Density Interval falling within the ROPE area. If all the HDI falls within the ROPE area (100%) the effects of the parameters can be considered negligible (Kruschke, 2014; McElreath, 2020). The closer the proportion inside the ROPE is to zero, the more confident are the effects of the parameters. Rhat is the ratio of the effective sample size to the overall number of iterations, with values close to one indicating convergence values.

| Class | System | Parameter | Median | CI (low) | CI (high) | ROPE (low) | ROPE (high) | %ROPE | Rhat |
| --- | --- | --- | --- | --- | --- | --- | --- | --- | --- |
| Amphibians | Freshwater | Intercept | 0.91 | 0.58 | 1.26 | -0.11 | 0.11 | 0.00 | 1.00 |
|  |  | Body mass | -0.15 | -0.47 | 0.22 | -0.11 | 0.11 | 0.36 | 1.00 |
|  | Terrestrial | Intercept | 1.44 | 0.97 | 1.93 | -0.14 | 0.14 | 0.00 | 1.00 |
|  |  | Body mass | -0.38 | -0.82 | 0.05 | -0.14 | 0.14 | 0.11 | 1.00 |
| Birds | Freshwater | Intercept | 1.06 | 0.92 | 1.22 | -0.11 | 0.11 | 0.00 | 1.00 |
|  |  | Body mass | 0.01 | -0.13 | 0.15 | -0.11 | 0.11 | 0.92 | 1.00 |
|  | Terrestrial | Intercept | 0.95 | 0.84 | 1.06 | -0.10 | 0.10 | 0.00 | 1.00 |
|  |  | Body mass | 0.29 | 0.18 | 0.40 | -0.10 | 0.10 | 0.00 | 1.00 |
|  | Marine | Intercept | 1.18 | 1.03 | 1.33 | -0.10 | 0.10 | 0.00 | 1.00 |
|  |  | Body mass | 0.16 | 0.02 | 0.30 | -0.10 | 0.10 | 0.18 | 1.00 |
| Bony Fish | Freshwater | Intercept | 1.12 | 0.98 | 1.26 | -0.08 | 0.08 | 0.00 | 1.00 |
|  |  | Body mass | 0.09 | 0.00 | 0.19 | -0.08 | 0.08 | 0.45 | 1.00 |
|  | Marine | Intercept | 0.77 | 0.71 | 0.83 | -0.07 | 0.07 | 0.00 | 1.00 |
|  |  | Body mass | 0.03 | -0.01 | 0.08 | -0.07 | 0.07 | 0.98 | 1.00 |
| Cartilaginous Fish | Marine | Intercept | 0.96 | 0.84 | 1.08 | -0.05 | 0.05 | 0.00 | 1.00 |
|  |  | Body mass | 0.09 | 0.01 | 0.16 | -0.05 | 0.05 | 0.16 | 1.00 |
| Mammals | Freshwater | Intercept | 1.14 | 0.65 | 1.62 | -0.12 | 0.12 | 0.00 | 1.00 |
|  |  | Body mass | 0.45 | 0.11 | 0.79 | -0.12 | 0.12 | 0.00 | 1.00 |
|  | Terrestrial | Intercept | 0.95 | 0.84 | 1.05 | -0.10 | 0.10 | 0.00 | 1.00 |
|  |  | Body mass | 0.30 | 0.19 | 0.39 | -0.10 | 0.10 | 0.00 | 1.00 |
|  | Marine | Intercept | 1.01 | 0.78 | 1.24 | -0.09 | 0.09 | 0.00 | 1.00 |
|  |  | Body mass | -0.04 | -0.26 | 0.19 | -0.09 | 0.09 | 0.58 | 1.00 |
| Reptiles | Freshwater | Intercept | 1.27 | 1.09 | 1.49 | -0.09 | 0.09 | 0.00 | 1.00 |
|  |  | Body mass | -0.01 | -0.21 | 0.20 | -0.09 | 0.09 | 0.63 | 1.00 |
|  | Terrestrial | Intercept | 0.69 | 0.10 | 1.34 | -0.07 | 0.07 | 0.00 | 4.97 |
|  |  | Body mass | -0.35 | -0.62 | 0.31 | -0.07 | 0.07 | 0.06 | 2.59 |
|  | Marine | Intercept | 1.52 | 1.15 | 1.86 | -0.09 | 0.09 | 0.00 | 1.00 |
|  |  | Body mass | -0.06 | -0.25 | 0.12 | -0.09 | 0.09 | 0.59 | 1.00 |

**Table S4. Outputs of the specific body mass models.** Median represents the median of the posterior distribution. CI low and high are the lower and higher values of the 95% confidence interval. ROPE represents the region of practical equivalence, which defines an area (lower and high 95% ROPE) where the value of the parameters is very likely to have a negligible effect (Kruschke, 2014; McElreath, 2020). %ROPE represents the proportion of the 89% Highest Density Interval falling within the ROPE area. If all the HDI falls within the ROPE area (100%) the effects of the parameters can be considered negligible (Kruschke, 2014; McElreath, 2020). The closer the proportion inside the ROPE is to zero, the more confident are the effects of the parameters. Rhat is the ratio of the effective sample size to the overall number of iterations, with values close to one indicating convergence values.

| Class | System | Parameter | Median | CI (low) | CI (high) | ROPE (low) | ROPE (high) | %ROPE | Rhat |
| --- | --- | --- | --- | --- | --- | --- | --- | --- | --- |
| Amphibians | Freshwater | Intercept | 1.08 | 0.72 | 1.44 | -0.12 | 0.12 | 0 | 1.00 |
|  |  | Latitude | 0.19 | -0.09 | 0.45 | -0.12 | 0.12 | 30 | 1.00 |
|  | Terrestrial | Intercept | 1.52 | 1.04 | 1.98 | -0.14 | 0.14 | 0 | 1.00 |
|  |  | Latitude | -0.03 | -0.43 | 0.40 | -0.14 | 0.14 | 52 | 1.00 |
| Birds | Freshwater | Intercept | 1.05 | 0.91 | 1.19 | -0.11 | 0.11 | 0 | 1.00 |
|  |  | Latitude | -0.09 | -0.19 | 0.02 | -0.11 | 0.11 | 67 | 1.00 |
|  | Terrestrial | Intercept | 0.82 | 0.72 | 0.92 | -0.10 | 0.10 | 0 | 1.00 |
|  |  | Latitude | -0.09 | -0.18 | 0.00 | -0.10 | 0.10 | 65 | 1.00 |
|  | Marine | Intercept | 1.10 | 0.95 | 1.24 | -0.10 | 0.10 | 0 | 1.00 |
|  |  | Latitude | -0.06 | -0.16 | 0.04 | -0.10 | 0.10 | 79 | 1.00 |
| Bony Fish | Freshwater | Intercept | 0.31 | 0.17 | 0.46 | -0.08 | 0.08 | 0 | 1.00 |
|  |  | Latitude | -0.69 | -0.73 | -0.65 | -0.08 | 0.08 | 0 | 1.00 |
|  | Marine | Intercept | 0.72 | 0.66 | 0.78 | -0.07 | 0.07 | 0 | 1.00 |
|  |  | Latitude | -0.04 | -0.07 | -0.01 | -0.07 | 0.07 | 100 | 1.00 |
| Cartilaginous Fish | Marine | Intercept | 0.89 | 0.79 | 0.98 | -0.05 | 0.05 | 0 | 1.00 |
|  |  | Latitude | 0.00 | -0.04 | 0.04 | -0.05 | 0.05 | 100 | 1.00 |
| Mammals | Freshwater | Intercept | 1.00 | 0.39 | 1.65 | -0.12 | 0.12 | 0 | 1.00 |
|  |  | Latitude | 0.09 | -0.31 | 0.48 | -0.12 | 0.12 | 43 | 1.00 |
|  | Terrestrial | Intercept | 0.81 | 0.70 | 0.91 | -0.10 | 0.10 | 0 | 1.00 |
|  |  | Latitude | -0.32 | -0.40 | -0.23 | -0.10 | 0.10 | 0 | 1.00 |
|  | Marine | Intercept | 0.99 | 0.76 | 1.23 | -0.09 | 0.09 | 0 | 1.00 |
|  |  | Latitude | -0.02 | -0.15 | 0.09 | -0.09 | 0.09 | 89 | 1.00 |
| Reptiles | Freshwater | Intercept | 1.02 | 0.37 | 1.73 | -0.07 | 0.07 | 0 | 1.00 |
|  |  | Latitude | 0.00 | -0.39 | 0.31 | -0.07 | 0.07 | 71 | 1.00 |
|  | Terrestrial | Intercept | 1.28 | 1.08 | 1.48 | -0.09 | 0.09 | 0 | 1.00 |
|  |  | Latitude | 0.13 | -0.07 | 0.31 | -0.09 | 0.09 | 32 | 1.00 |
|  | Marine | Intercept | 1.53 | 1.24 | 1.83 | -0.09 | 0.09 | 0 | 1.00 |
|  |  | Latitude | -0.13 | -0.29 | 0.01 | -0.09 | 0.09 | 27 | 1.00 |

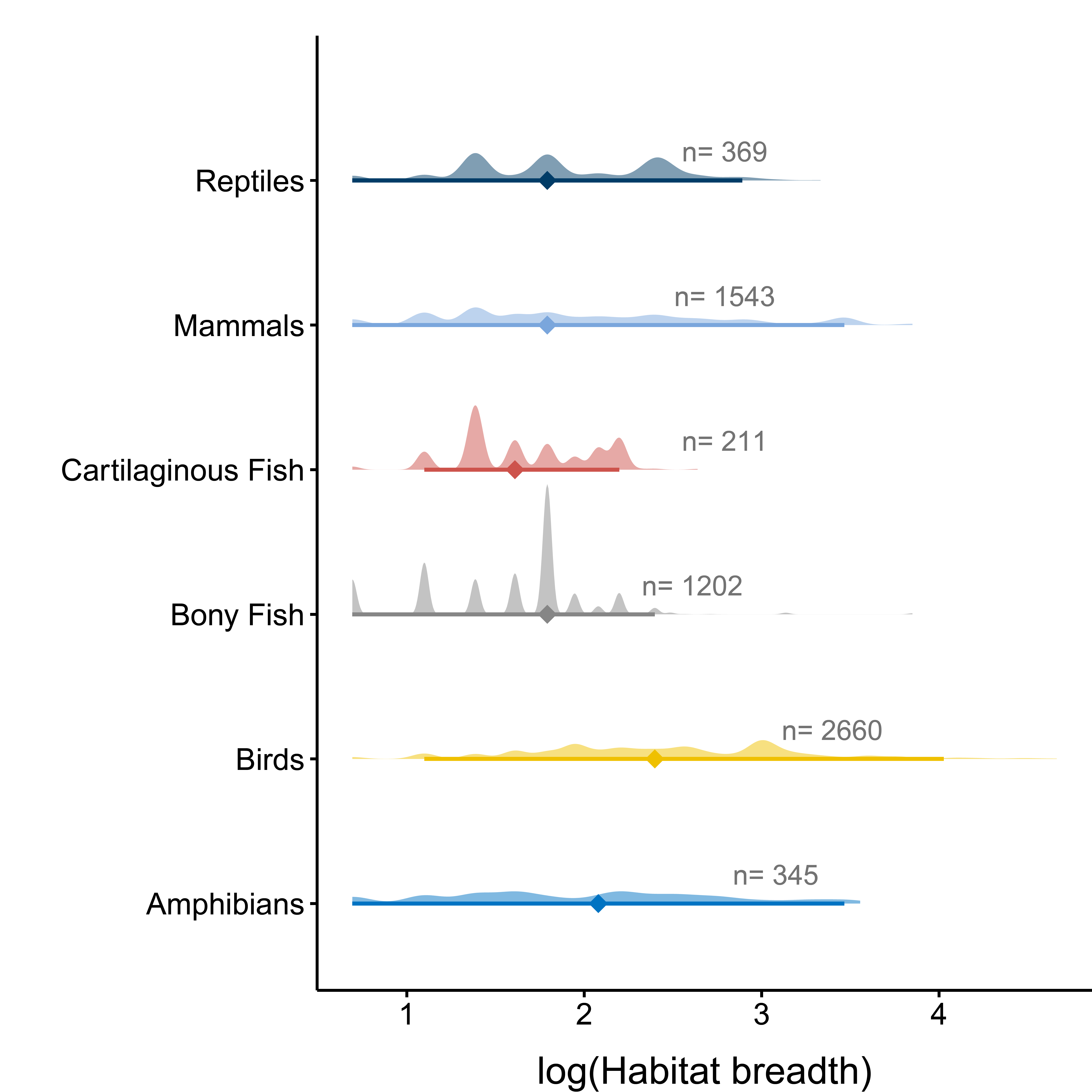

**Fig. S2. Habitat breadth across the different taxonomic classes.** Distribution of the raw data of habitat breadth (natural logarithm transformed) across the taxonomic classes of the species included in this study. The lines below the distribution represent the 95% confidence intervals, with the dot representing the median value. n values represent the sample size.

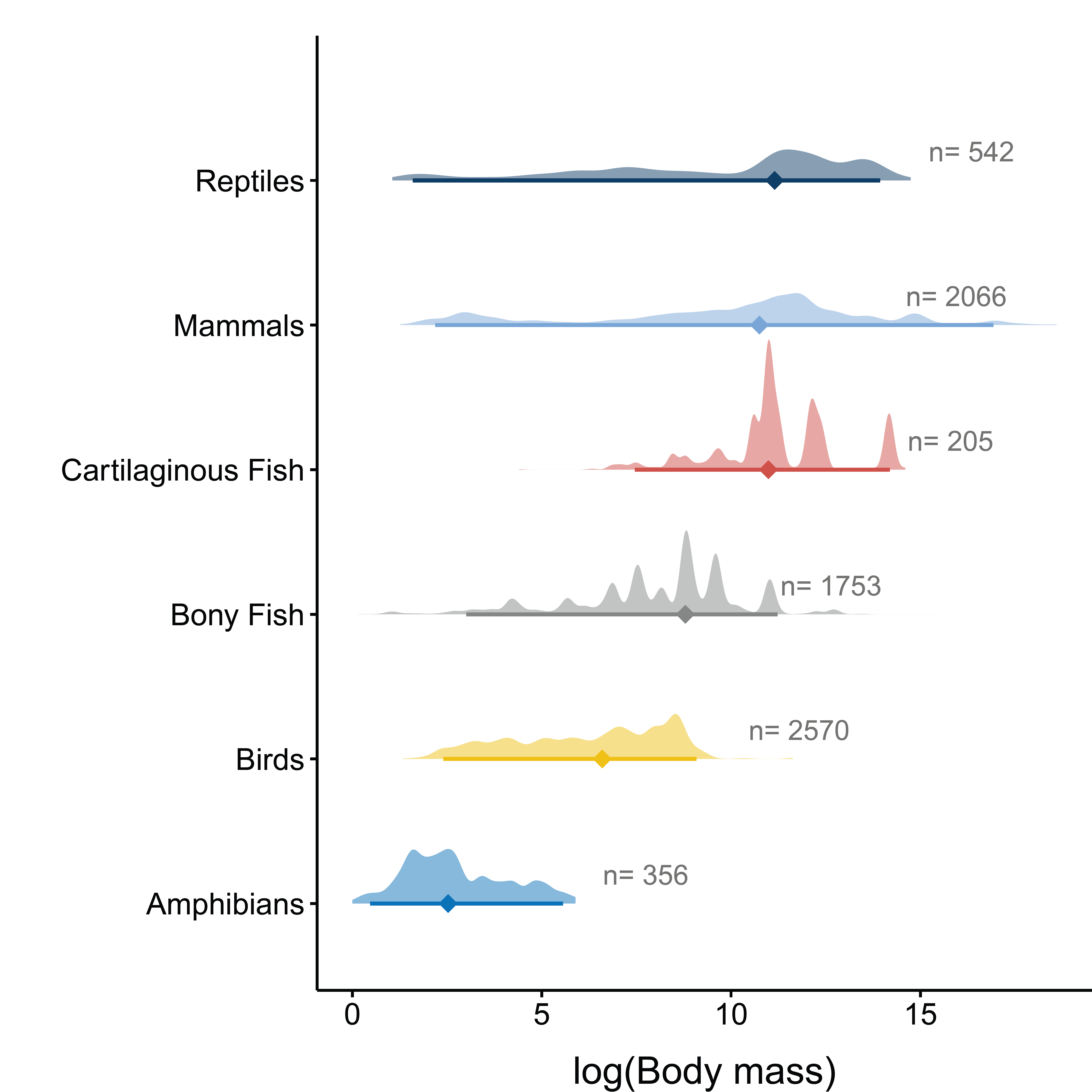

**Fig. S3. Body mass distribution across the different taxonomic classes.** Distribution of the raw data of body mass (natural logarithm transformed) across the taxonomic classes of the species included in this study. The lines below the distribution represent the 95% confidence intervals, with the dot representing the median value. n values represent the sample size.
